# HLAssign 2.0: An advanced Graphical User Interface for the analysis of short and long read Human Leukocyte Antigen-typing data

**DOI:** 10.1101/2020.05.25.087627

**Authors:** M Wittig, M Schmöhl, S Koch, M Ziemann, S Görg, V Lange, F Jacob, M Forster, A Franke

**Affiliations:** Institute of Clinical Molecular Biology, Christian-Albrechts-University of Kiel, 24105 Kiel, Germany; Muthesius University of Fine Arts and Design, Interface Design, 24103 Kiel, Germany; Institute for Transfusion Medicine, UKSH Campus Lübeck, 23562 Lübeck, Germany; Institute for Transfusion Medicine, UKSH Campus Kiel, 24105 Kiel, Germany; DKMS Life Science Lab, 01307 Dresden, Germany

## Abstract

Next Generation Sequencing (NGS) based Human Leukocyte Antigen (HLA) typing has been a challenge due to the polymorphism of the HLA region. Nevertheless, the method’s accuracy has increased during the last years and it is now routinely used by many large centers including bone marrow registries. However, challenging HLA genotype compositions exist, which hinder a fully automated analysis. Therefore, HLA typing results are still visually inspected in diagnostics, i.e. the underlying read mappings and phasing information is controlled. Here, we present HLAssign 2.0 that now includes a strict workflow, improved tools for visual inspection and read phasing analysis in the automatic caller. In collaboration with interface design researchers, biologists and informaticians we developed an elaborate graphical user interface for visual evaluation of automated HLA calls for Illumina NGS reads. We also provide tools to preprocess 10x Genomics and PacBio sequencing reads for HLAssign analysis. We benchmarked our automatic caller against STC-seq and xHLA, showing comparable automatic call rates. Additional manual inspection of the automatic results in our GUI assists the user to assign the correct HLA calls and to achieve diagnostic accuracy. HLAssign 2.0 is free for research and commercial use and is available for Windows and MacOS.

## Introduction

The Human Leukocyte Antigen (HLA) complex is a highly polymorphic region of chromosome 6 and plays a major role in the adaptive immune system. There is rapidly growing interest in studying HLA associations in various diseases^1,2^. Next Generation Sequencing (NGS-) based HLA typing approaches were developed in the last years and are beginning to replace classical typing methods especially in laboratories with a higher sample throughput and the need of a higher resolution^3–7^. Bioinformatic typing algorithms range from simple read count analyses per hypothetical allele, that may include complex filtering criteria, that sometimes employ a *de novo* assembly with back mapping to known references, up to iterative matching and filtering steps^4,6–9^. Nevertheless, unambiguous HLA typing remains a challenge as recombination of the underlying polymorphisms often leads to multiple equally possible genotype calls^10^. Paired end sequencing of both ends of a DNA fragment allows for some limited phasing between the read pairs, which can improve the HLA calling of “difficult genotypes”. The limitation depends on the SNP density of the underlying HLA genotypes and the paired end distances of the sequenced reads, because at least two SNPs have to be covered by such a single or paired read. The paired end distance is limited by the density of the sequencer clustering and preferential amplification of shorter fragments. A mixed library that includes both paired end and mate pair fragments would achieve a wider range of paired read distances and thus allow for phasing over a longer genomic distance. Nevertheless, challenging genotypes that cannot be phased will remain. The most recent long read sequencing technologies, e.g. from Pacific Biosciences or Oxford Nanopore ^11,12^, may further improve the phasing and calling as the genomic variation is captured on a single molecule that stretches over several kilobases. Other technologies – e.g. from 10x Genomics – utilize an emulsion-based library preparation step, in which longer DNA fragments from a single chromosome are captured in a droplet – and therefore a larger physical haploid DNA segment – and are fragmented into sequencing reads that are tagged with the same unique molecular barcode^13^. The 10x method allows for phasing of multiple reads or fragments that originate from a single DNA molecule which may yield highly accurate and unambiguous HLA typing results. However, a visual inspection of the read mappings for a specific genotype is still the gold standard in diagnostic routine as this also helps to identify false positive HLA calls and resolve ambiguous allele calls. To the best of our knowledge, no open-source and non-commercial HLA typing software tool currently exists that offers a graphical user interface (GUI). This may be due to the circumstance that bioinformaticians often focus on the algorithmic development, prefer command line tools and may not be that familiar with time-consuming GUI-development. We here present a successor version of our previously reported HLAssign tool(6). The new version 2.0 includes a strict and intuitive workflow, comes with a significantly improved GUI, and can include phasing information. The GUI and the analysis workflow were iteratively developed by professional interface design researchers in close interaction with HLA typing laboratories.

Since our publication of HLAssign 1.0, several new HLA calling tools for NGS data have been published. The most recent publications about free-for-academic-use tools are those about xHLA^7^ and STC-seq^8^ – to our knowledge these latter tools are also the currently best performing free tools. xHLA iteratively refines the mapping results at the amino acid level to achieve four-digit HLA types. STC-seq evaluates the equal mapping across all known HLA loci to identify the best match for the underlying NGS data. Even though these new tools perform slightly better than older tools, they do not work perfectly, i.e. achieve >99.9% diagnostic accuracy. Therefore, HLAssign 2.0 can help in the analysis of difficult samples by providing visual inspection and options for manual correction of the automated HLA calling.

## Results and Discussion

To assist users in manual review of HLA typing data, we implemented several new features in HLAssign 2.0, in an iterative development process which involved user testing and iterative implementation of features that addressed user feedback. The final GUI that we present here thus implements features to support the decision-making processes that we learned from HLA typing experts in the four typing laboratories that we visited (see Acknowledgements). Version 2.0 is therefore a significant improvement compared to the simple GUI of Version 1.0, which was developed by a bioinformatician alone (for comparison, see Figures 1 & 2). The new interface is not only visually more sophisticated, it also assists the user to do a more efficient and accurate analysis based on expert processes in the HLA typing laboratories.

**Figure 1.**
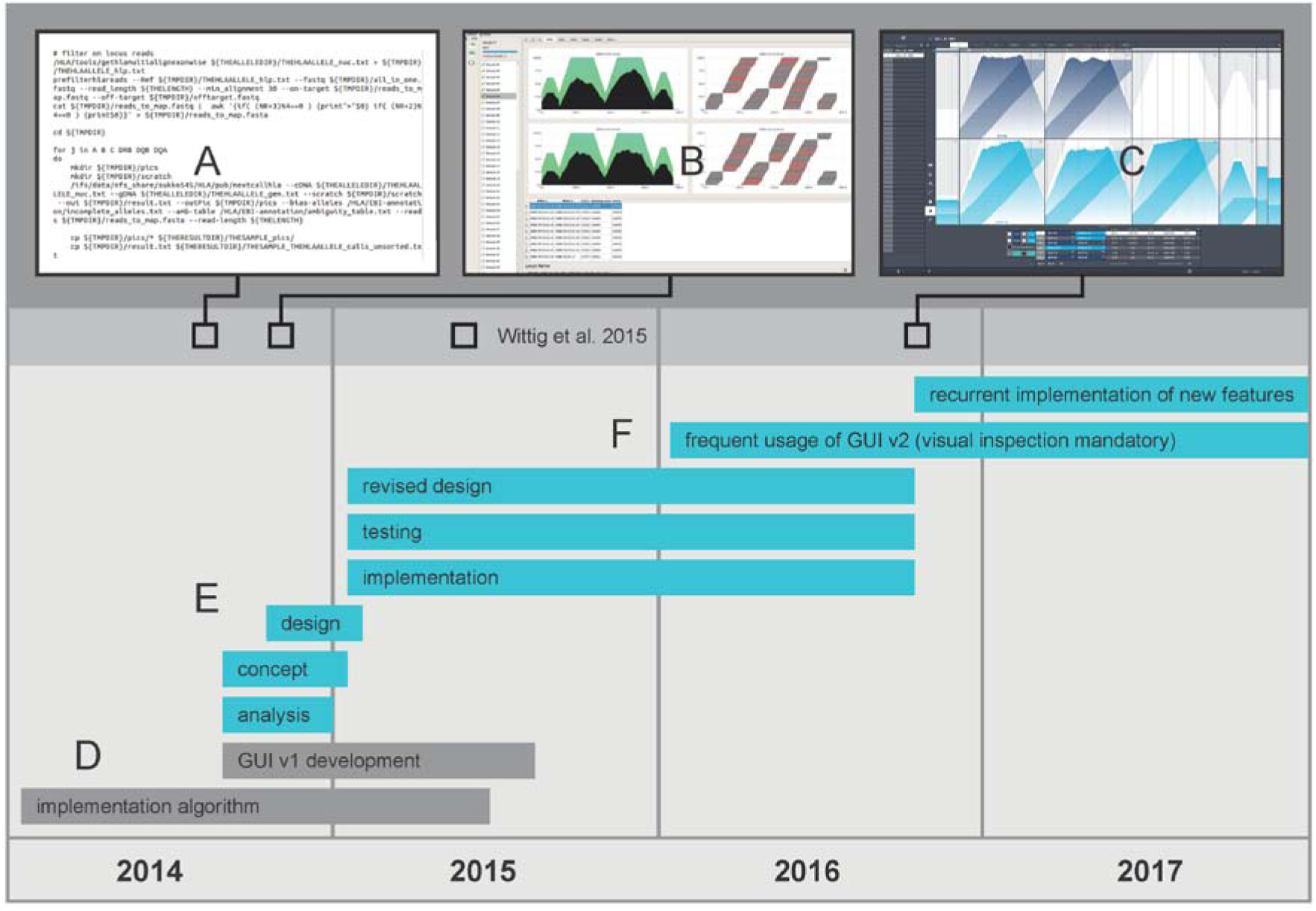
GUI development workflow: The workflow shown above illustrates the different phases and tasks that were performed during the development. **(A)** Terminal application with a text-based user interface at the start of the project. **(B)** First graphical user interface (GUI) prototype in 2014. **(C)** Intermediate GUI v2 prototype in 2016 before phasing was added. **(D)** The grey bars show the start of the project with the development of the analysis algorithm, finally published in June 2015. **(E)** The turquoise bars of the workflow visualize the tasks performed for the GUI v2 development. The analysis and concept development were parallel processes and after these tasks showed first progress we were able to start with the conceptual designs. These initial tasks were finished after only a couple of months and we could start implementing the design ideas. The implementation phase is dominated by an intensive communication and interaction of designer, programmer and user. Design ideas were changed and re-implemented when practical user cases required solutions that differed from the initial idea. **(F)** From 2016 onwards, we used the GUI v2 frequently and visually inspected all HLA calls. This routine usage brought up ideas of useful new features, such as visualization of cross locus mappings or cis-SNPs and their incorporation into the automatic caller, or a third locus mapping display track to enable the comparison of mappings with alternative alleles. These new features were each implemented in the course of software updates.

### Role model

In addition to presenting our new software version, we here also present a role model for how a professional GUI development can be driven in the academic setting. As reported here, it helps in practice if such professional-level GUI developments are performed by an interdisciplinary team of experts. Software tools that may benefit from the herein-described GUI development workflow include for example those tools that are frequently used by the community and that are still maintained.

### GUI and workflow design

In HLAssign 2.0, besides improving the interface design, several improvements were made in the data analysis workflow, which is described in the following. Firstly, the user is guided through an expert workflow when submitting samples for analysis (**Supplement S1**). This assists the user to process a sample with all necessary steps and to assign necessary meta-data and analysis settings. We also eliminated the need to use multiple software tools. For example, at the beginning of the analysis process, the user has the option to display the quality metrics of the selected raw sequencing files (in fastq format) and to adjust sequence trimming based on this information, without the necessity of using extra tools such as FastQC and Trimmomatic. We redesigned the data display (Figure 2) based on an information value concept that provides and ranks only the important information. The genotype table in the data display contains the relevant calling parameters. High scores in each parameter and the frequency of occurrence of each allele (“common defined (c)”, “well defined (wd)”) are highlighted according to the information value concept. Less relevant parameters are hidden and not shown to the user. Incompletely covered alleles and alleles failing quality control are available in a dialog window and not shown directly (“benefit analysis”). Each locus has a color highlighting the estimated quality of the analysis. Because the system lacks self-explaining decision making and verification, the user has to be trained in the importance and consequences of each parameter, because they are related to each other and affect the decision making process. The graphical visualization of the result is in general more important than the parameters and therefore more prominent on the screen. Our user tests with an eye-tracking software showed that the experienced user focuses first on the graphical visualization and if any anomalies appear, the user evaluates the heterozygous nucleotides of a genotype and finally the parameters in the table. Although the user has to learn several interrelated variables of the information values, the expert decision-making process can now be learned and performed more confidently. Usability tests of the previous software version 1.0 showed that users who are not familiar with the calling algorithm (even though the important metrics are described in the tutorial and in Wittig *et al.* 2015^6^) rely on the score and erroneously decide that the result is highly confident if the score is high. The score in version 1.0 was the harmonic mean of the collected mapping metrics, which was aimed at providing a clue but was not aimed at reflecting the most confident call. This example illustrates how a GUI may confuse a user if it was designed by programmers and close colleagues without any independent external user feedback. After realizing that the users rely on a score rather than on a tutorial or algorithm description, we changed the way that the score is computed: It is important for a confident call that the best score is significantly higher than the second-best score. Therefore, in version 2.0 we replaced the old score by a penalty that represents the difference to the highest score.

**Figure 2.**
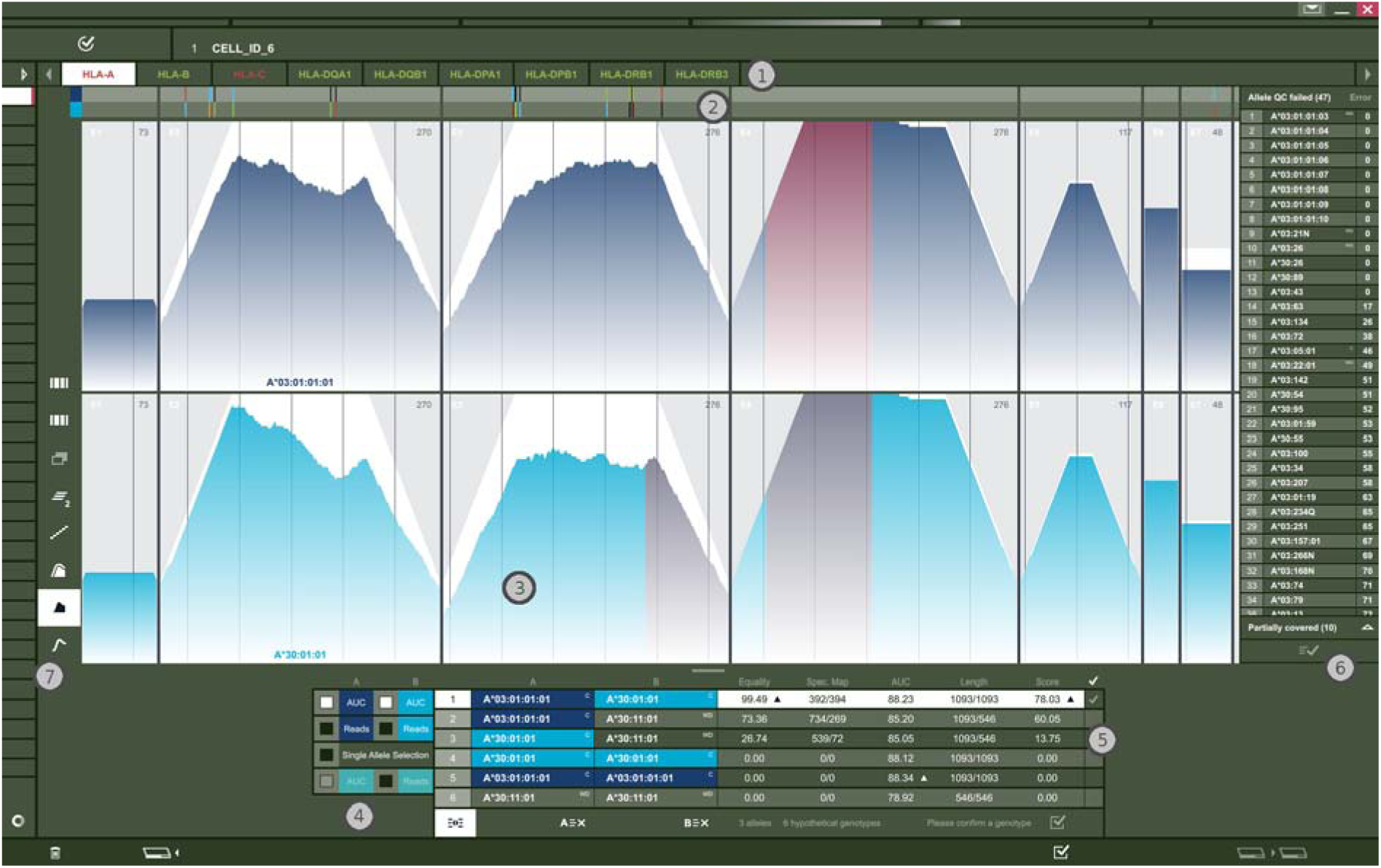
Data Visualization in HLAssign 2.0: The figure shows the *validation display* which is the main interactive tool for visualizing and validating the automated calling, removing alternative alleles and assigning the correct HLA calls. To perform an accurate validation, it is important to understand the characteristics of the underlying data. The most important point to consider is the random fragmentation that is normally performed for all samples prior to sequencing. This means that we expect a nearly even distribution of the mapped reads across the entire locus and both alleles of a given genotype. The more balanced these distributions are, the more confident the calling of the genotype will be. The different sections of the validation display are described in the following numbered panels: **(1)** Upper menu bar selects the tabs for the different loci. **(2)** Consensus bar shows all variations (SNVs) of the currently displayed genotype. **(3)** Read mapping windows show the raw data for each allele of a genotype at each locus. **(4)** Check boxes that toggle between different raw data views. **(5)** Table that shows the ranked genotypes and that is sorted by the penalty calculated by the automated caller. **(6)** Lists of automatically filtered (rejected) alleles. Here, the user can switch between two lists. The first list contains all alleles that are completely covered by reads but filtered out by the first filtering step (unbalanced read distribution). The second list contains some of the alleles that are not completely covered by mapped reads; it is sorted by uncovered bases in ascending order. **(7)** Buttons to change the visualization style. A detailed description can be found in **Supplement S1** (Supplementary Figure 8).

### Resolution of ambiguities

Ambiguities in HLA calling are often caused by homologous sequences at other loci or by the high similarity of different alleles of the same gene. An example for homologous sequences at other loci is HLA-A*32:101Q which has the same exon 3 sequence as HLA-H*02:01. The remaining HLA-A*32:101Q sequence equals HLA-A*32:01. So if HLA-A*32:01 and HLA-H*02:01 are present in the same sample, false positive signals may occur, especially when targeted capturing was performed (as described in ^6^). To warn the user of this sample-dependent problem, HLAssign highlights the respective coverage and/or reads in red if such an event occurs. The second issue, high similarity of different alleles of the same locus can be found very frequently. This may result in the phenomenon that *in-silico* recombination of stretches of a given genotype may lead to plausible but wrong genotype calls (the typical Sanger based sequencing issue when no allele specific sequencing primers are used). This issue can be solved by determining haplotypes that are as long as possible. The longer the haplotype is, the less recombination ambiguities arise and the more confident the genotype call is. In HLAssign 2.0, the new visual feature that highlights SNPs from the same haplotype helps to detect these cases. This allows for highly confident data curation of both described phenomena. The new software release also allows for the visual inspection of three alleles simultaneously. This makes it possible to select a putative genotype and add an alternative allele to observe the underlying mapping in the same context (**Supplement S2**). In addition, to make use of the benefit of long haplotypes we released two auxiliary programs that convert Pacific Biosciences and 10x Genomics fastq files for usage in HLAssign. These programs can be downloaded from GitHub (https://github.com/MiWitt/10xToFastq, https://github.com/MiWitt/FragmentLongReads).

### Benchmark of automatic callers

We validated accuracy with a reference data set from regular blood donors of northern Germany. The reference HLA genotypes were generated with the method described by Lange *et al.*^14^ and have G-group resolution. This method was already used for millions of samples and as it is used in routine bone marrow donor matching it is highly validated. Figure 3 shows the concordance rates between the reference set for HLAssign, xHLA and STC-seq. All tools show comparable concordance rates. As xHLA delivers 2 field resolution only, we always assumed the reference G-group if the assignment was ambiguous. STC-seq and HLAssign deliver 3 field resolution and unambiguous G-group assignment was possible. The full benchmark data set is provided as **Supplement S3**. For HLAssign v1, Jiao *et al*.^8^ showed that the automated calling performed best for the old reference data set but that it loses accuracy when applied to other data. We observed a similar effect. It turned out that the old reference data set – that was optimized for diversity and by random permutations assembled with a minimal sample number – was not reflecting realistic populations where problematic genotypes can occur multiple times. To validate our algorithm, we switched to the completely new data set presented in Figure 3 and could show that the automatic caller in HLAssign v2 was improved.

**Figure 3.**
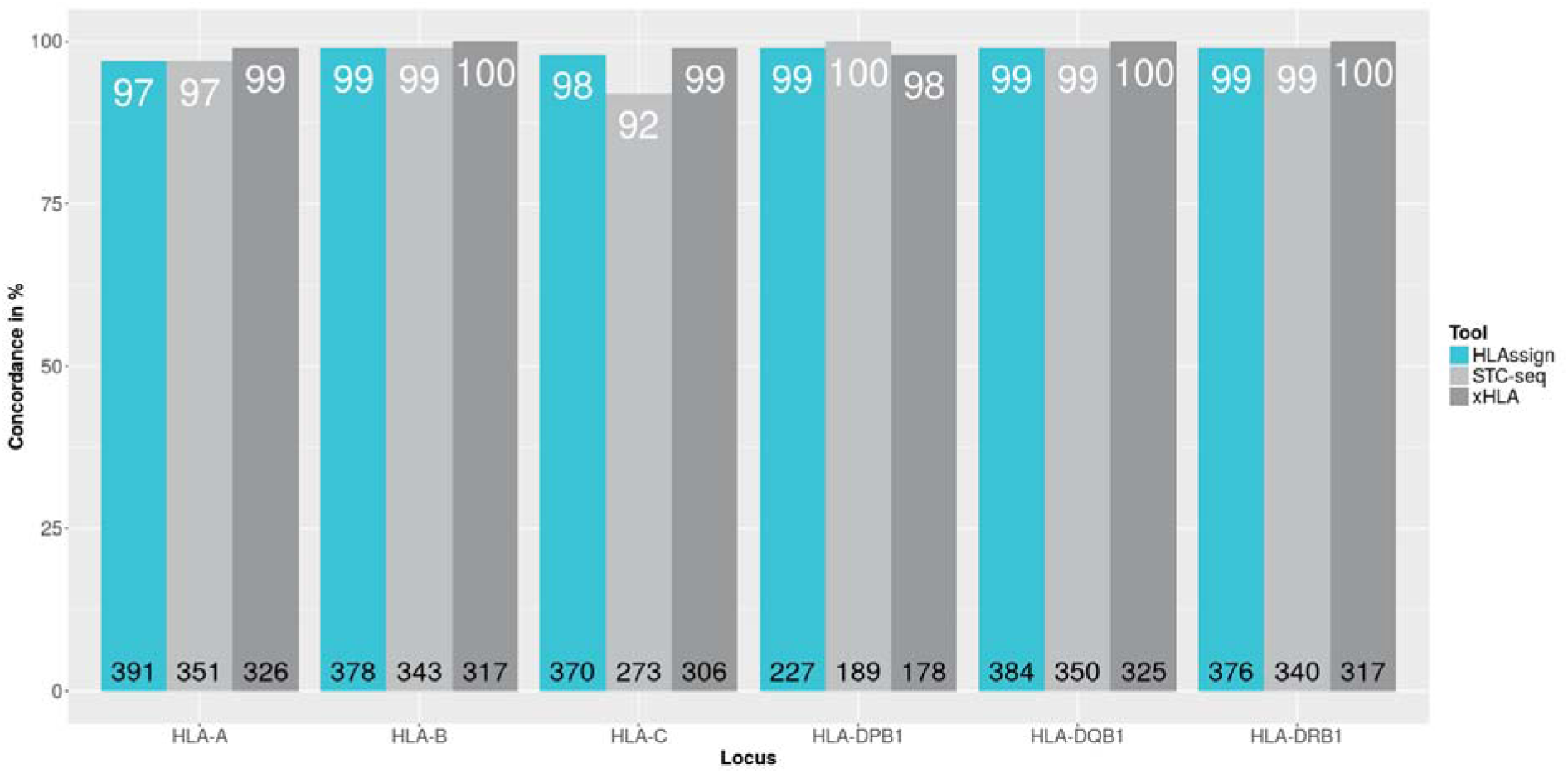
Concordance rates of automated analysis: This figure shows the concordance in percent of the new HLAssign 2.0, STC-seq and xHLA compared to the reference set (DKMS(14)). The black numbers at the bottom of each bar group show the number of available genotypes for that locus. Available genotypes are genotypes with at least 4-digit unambiguous calls in the reference set and with genotype calls in all data sets (reference, HLAssign, STC-seq and xHLA). The white numbers correspond to the bar height and show the concordance to the reference data set in percent. Overall there are 170 unique HLA-A alleles forming the 391 different HLA-A genotypes of panel 1. For the other loci there are 170 unique HLA-B alleles, 170 HLA-C, 12 HLA-DPB1, 16 HLA-DQB1 and 32 HLA-DRB1. Accuracies are near 100% after manual review within the new HLAssign 2.0 GUI software.

### Conclusion

In conclusion, the aim of our second major software version was to improve the robustness of the automated calling and to assist researchers in evaluating the data visually. In Figure 3 we show that the new algorithm performs comparably to recently published tools. As the benchmark data set is completely new and neither our algorithm nor those of xHLA or STC-seq were optimized for it^8^, we could show that our new algorithm performs well on an arbitrary data set. Nevertheless, there are still errors in the automatic typing, but these may also occur using the xHLA and STC-seq algorithms (**Supplement S2**, **Supplement S3**). In summary, our benchmark shows that the NGS-based HLA calling accuracy of academic tools has increased during the last years, but that they are not perfect. However, with visual inspection and guided data evaluation, a user can generate almost perfect results that meet diagnostic accuracy.

## Material and Methods

### Benchmark samples

Benchmarking samples were collected in 2014 by the Institute of Transfusion Medicine in Lübeck, from 434 regular healthy blood donors of North German ancestry. All donors included in the study gave written informed consent for research use. The study was approved by the ethics committees of the Christian-Albrechts-Universität zu Kiel (ID: A103/14) and all subjects provided written informed consent. The applied methods include DNA extraction, DNA amplification, NGS and data analysis, and are described in the next section.

### NGS and HLA-Typing for the benchmark

All benchmarking samples were HLA-typed at G-Group resolution by the DKMS Life Science lab in Dresden^14^. The DKMS laboratory performs HLA typing for allogeneic stem cell transplantation, operating one of the world’s highest-throughput HLA typing platforms. Hence the DKMS HLA typing results count as the current gold standard for HLA typing. In parallel, the samples were sequenced at our institute, the Institute of Clinical Molecular Biology, with our previously published protocol^6^ on a HiSeq 2500 with 2×100bp paired-end reads and 96 samples per lane. Typing with xHLA^7^ and STC-seq^8^ was performed with default settings. For benchmarking the automated callers, analysis with HLAssign 2.0 was performed without any manual corrections using the GUI. Concordance rate was calculated for all samples and loci where all tools (xHLA, STC-seq and HLAssign) called a genotype and where the reference genotype was unambiguous (no NMDP code).

### GUI

The new user interface was implemented in Qt Meta-object Language (QML). QML is a declarative language that allows user interfaces to be described in terms of their visual components and how they interact and relate with one another. QML is a highly readable language that was designed to enable components to be interconnected in a dynamic manner. The QML user interface is connected to a back-end C++ library for the HLA typing algorithm. To interconnect QML and the core algorithm, a C++ API based on Qt (pronounced “cute”) was created (www.qt.io). Qt is a cross-platform application framework that is widely used for developing application software that can be run on various software and hardware platforms with little or no change in the underlying code base, while having the power and speed of native applications. QML and Qt code can be compiled/executed on all common operating systems such as Windows, Mac, Linux, Android and the same applies for C++. This allows for high portability of the software tool.

### Hardware

HLAssign analysis was performed on a 2.6 GHZ quad core and 24 GB RAM desktop computer. The recommended minimal memory and storage requirements are 8 GB RAM and 1 GB free disk space. xHLA and STC-seq analysis was performed on a computer cluster of AMD 3.6 GHz processors, assigning 24GB of RAM and two cores per sample.

### Haplotype estimation (phasing)

The new version considers phasing information for the automated calling algorithm. It performs a multiple alignment of all putative alleles and observes the read mappings at the SNP positions that discriminate between the different putative haplotypes. If two SNPs are covered by a single or paired read we call these SNPs *linked SNPs* (SNPs in cis). Our algorithm counts the number of links between discriminating SNP pairs, the number of linked SNPs and the number of mapped reads that phase two or more SNPs. Eventually it scores the genotype using these three measures and the old score^6^ and calculates an equally weighted harmonic mean of these four values.

### Design

The human-computer-interface design can heavily assist or influence a decision made by the user.^15^ For instance: The verification of the algorithmic HLA analysis depends only on the given information represented on the screen. In other words, the user decision is based on symbols (such as numbers or icons), graphical shapes, colors and the provided internal structure. To improve usability, the software must be converted into a decision supporting system (DSS). DSSs are well known in economic science where decisions are highly complex and depend on a huge amount of data.^16^ For our bioinformatics development project, we summarize this process in Figure 1. The method that we applied was the user centered design (UCD)^17,18^, for which three different types of specific knowledge are needed: (1) Knowledge about the technology. (2) Knowledge about the user, the user’s communication and interactions, and graphical design. (3) Knowledge about the task that has to be accomplished by the user. In highly specialized research projects, it is particularly vital to engage experts from the disciplines^17,19^. To achieve that knowledge the analysis phase started with on-site visits to four different HLA typing facilities where interviews and evaluation of the status quo were performed. With these insights and the knowledge of the new kind of data used for the NGS based HLA typing a conceptual design was created followed by the implementation of the design. During the implementation it was important to have a close interaction between designer, programmer and user. Feedback was essential to adapt the GUI to the needs of the user and to prioritize and weigh the important information accordingly.

### Software and data availability

The software, example data, full disclaimer and tutorial material are freely available without any registration requirements on the website http://www.hlassign.org

In brief, HLAssign 2.0 can be used free of charge under the condition that the user is solely responsible for the results and that this publication and the website are cited.

## Supporting information

Supplement S1, Workflow

Supplement S2, Examples data evaluation

Supplement S3, Samples and HLA calls

## Acknowledgements

We thank all laboratories and their staff for giving us the opportunity to gain insights into their daily HLA typing routine. These laboratories are: MVZ Dr. Eberhard & Partner Dortmund GbR (ÜBAG), Brauhausstraße 4, 44137 Dortmund, Germany; Institute of Clinical Medicine, Kirkeveien 166, Laboratoriebygget, 0450 Oslo, Norway; University of Lübeck, Institute of Transfusion Medicine, Lübeck, Germany.; Med. VersorgungszentrumMVZ DRK-Blutspendedienst Ulm gGmbH, Helmholtzstr. 10, 89081 Ulm, Germany.

## Author Contribution Statement

M.W. programmed the algorithm, performed the benchmark, worked on the GUI, prepared the figures and wrote the manuscript. M.S. implemented/programmed the GUI. S.K. designed the GUI. M.Z. and S.G. reviewed and corrected the manuscript. V.L. provided the reference data and reviewed the manuscript. F.J. supervised the GUI design. M.F. wrote the manuscript. A.F. supervised the project and wrote the manuscript.

