## Supplement S1, Workflow for "HLAssign 2.0: An advanced Graphical User Interface for the analysis of short and long read Human Leukocyte Antigen-typing data"

**HLAssign Workflow**

The graphical user interface (GUI) mainly consists of the workflow a typical sample goes through. The workflow steps are in this direction: Import, Identification, Locus, Calculation, Verification and Report. The different workflow views can be selected by the user by clicking the toolbar, which becomes visible by mouse over at the top of the application window. Figure 1 shows the afore-described toolbar. The main idea for the workflow is to assist the user with the sample analysis. It starts at the very left with the data import and ends at the very right with the report export. In every workflow step properties have to be assigned or HLA calls have to be validated/corrected and/or confirmed. If done, the sample status changes and it can be forwarded to the next workflow step. An action bar is located at the bottom of the application window (Figure 2). The action bar has buttons to perform dialog specific actions like, sample import, property assignment or sample forwarding.

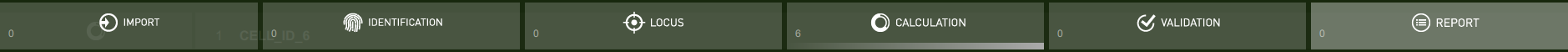

Figure 1. The toolbar of the application allows the user to select the different workflow steps, from left to right: Import, Identification, Locus, Calculation, Verification and Report. The currently selected view is highlighted in light-grey, which is Report in this example.

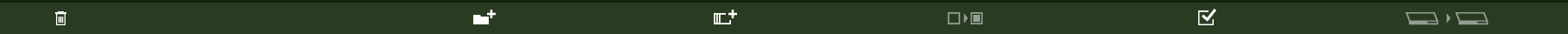

Figure 2. The action bar of the application, located at the bottom of the window. This action bar is from the import dialog, but 4 of its buttons can be found at every dialog. The very left button is the trash button. The next two buttons are dialog specific and are for importing multiple samples or a single sample. The fourth button is for assigning properties to multiple samples. The check button right of it is to confirm current settings for the currently selected sample. The very right button is active if at least one sample has confirmed settings. If pushed, it moves all samples with confirmed settings to the next workflow step.

In every view where properties like read trimming, project title, loci to analyze et cetera can be assigned, the user has the possibility to use a multi confirm functionality. This functionality allows assignment of current setting to multiple samples by doing only a few clicks. To do so, the user has to activate the multi confirm checkbox in the main view next to the current settings (Figure 3). If this box is checked, the multi confirm button in the action bar gets activated (fourth button in figure 2). If clicked, a dialog pops up and the user can select all the samples to which he would like to assign the currently made settings. If done, all these samples will get a light grey status bar and can be forwarded to the next workflow step. The samples can be found in a list aligned left to the main window. Each list entry consists of the order number followed by sample name and by a right-aligned status bar. This status bar is not visible if action, like read trimming, has to be assigned and becomes visible and grey if a sample is ready to be forwarded to the next workflow step. In the validation view a red status bar indicates samples where manual review is strongly recommended and a green status bar for samples where the automatic calling delivers robust results.

The first workflow step is the Import (Figure 4). It allows the user to add samples to the analysis and to perform quality inspection together with some read trimming functionality. Like all views, the import view consists of 4 areas. At the left side the user finds the sample list of the currently selected workflow step. Status information for the current workflow step can be found at the top, together with the workflow toolbar. At the bottom of the application are function to perform submit, forward- or reverse-operations, load files, et cetera. The remaining area, which covers the majority of the screen, is reserved for the workflow specific control elements and data views. In almost every view the user can find the multi confirm check box. If properties should be assigned to multiple sample, the multi-confirm buttons enables this functionality by enabling the multi confirm button at the action bar. When clicked, a modular dialog pops up and all samples that should get the selected property/properties value/s can be selected. This is especially useful to prevent dozens of separate confirms to a bulk of samples that have to be analyzed with the same settings.

The different workflow steps are in the following described by screenshots:

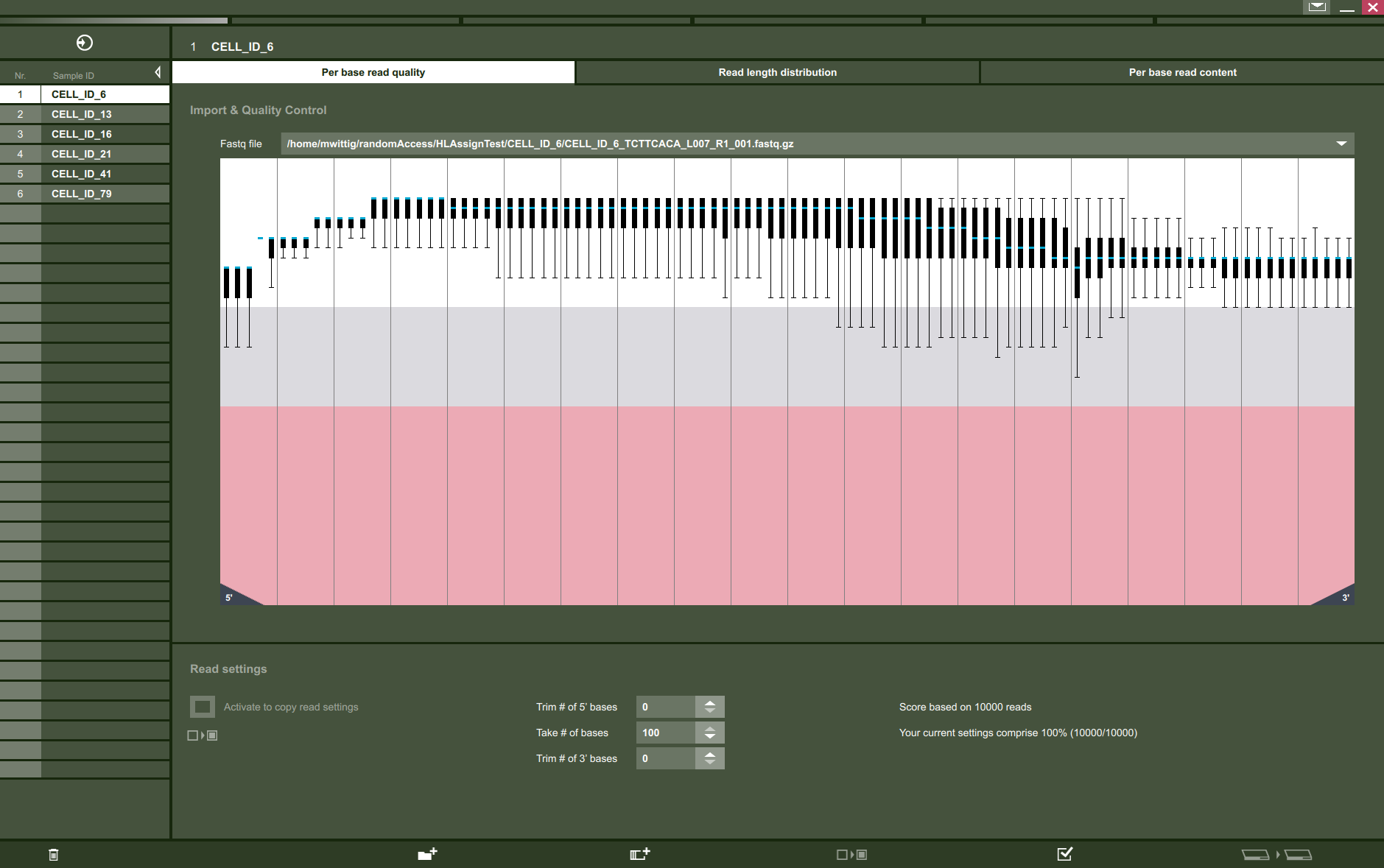

Figure 4. **Workflow Step 1**, the import. Choose *add directory* or *add files* to add fastq files of one or more samples. If selecting *add files*, the user can browse for 1-n fastq files and add them to a sample. The sample name can be given in the sub dialog that pops up. If selecting *add directory*, the user can select 1-n directories. The directory name is then taken as sample name and all fastq files within that directory are assigned to that sample. After the samples/fastq files are added, the first 10,000 reads of all files are processed to perform an initial QC. The QC consists of per base quality (main view above), read length distribution (in most cases one length for all reads) and per base read content. Based on that information the user can perform 5’ and 3’ read trimming. The chosen settings can be confirmed by the check button at the lower right (second from right). If the same settings should be applied for multiple samples, the user can use the button *activate to copy read settings* and push the *multi confirm* button left of the *confirm* button. This multi confirm functionality can be found in all the other workflow steps as well. With the *push next* button at the lower right, all confirmed samples are pushed to the next dialog. An animation starts to show explicitly that the samples are now available in the next step.

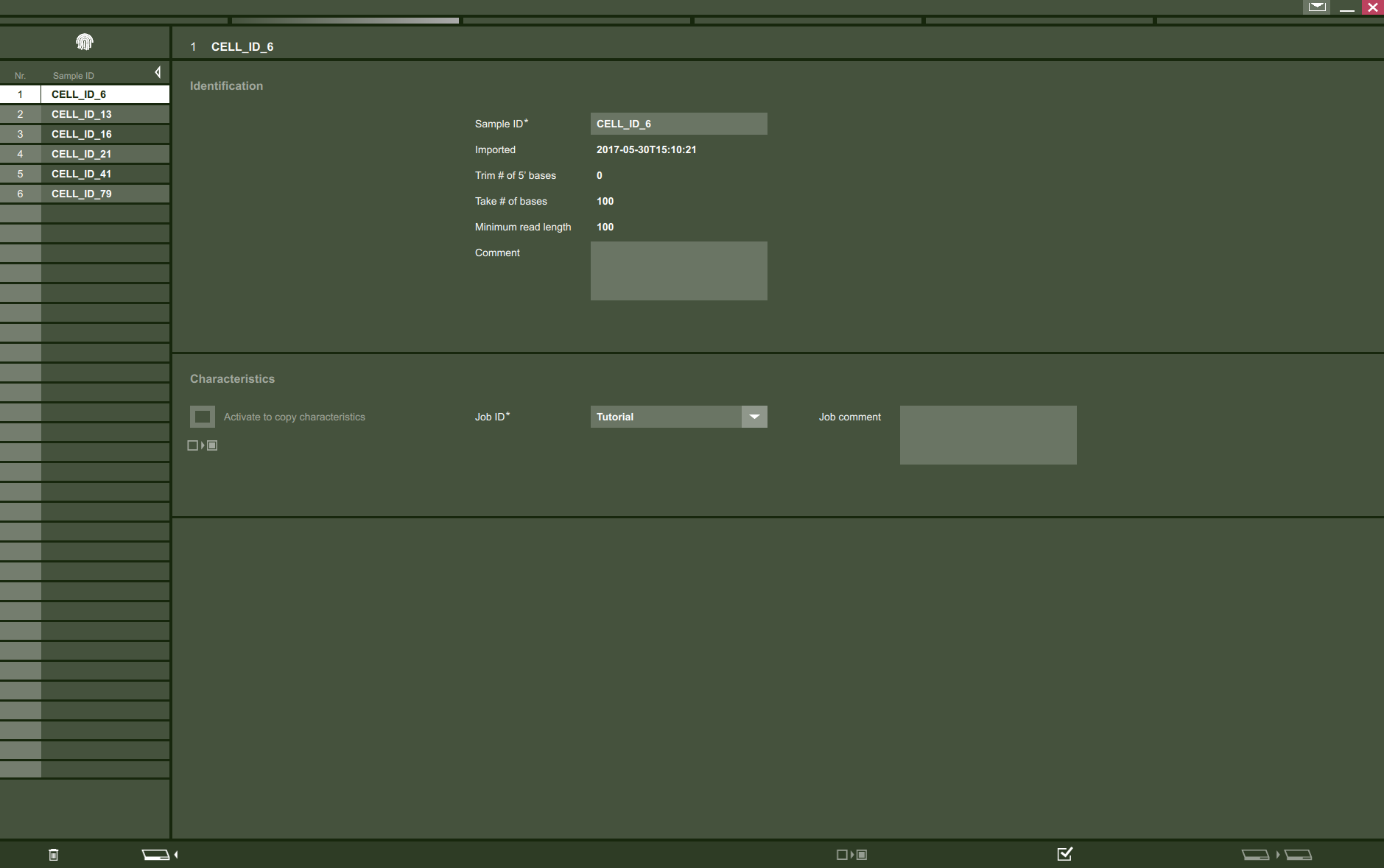

Figure 5. **Workflow Step 2**, identification. In this step, the user can change sample names, assign comments and a job Id. Similar to the previous workflow step the user can assign the job id setting to multiple samples instead of confirming each sample separate.

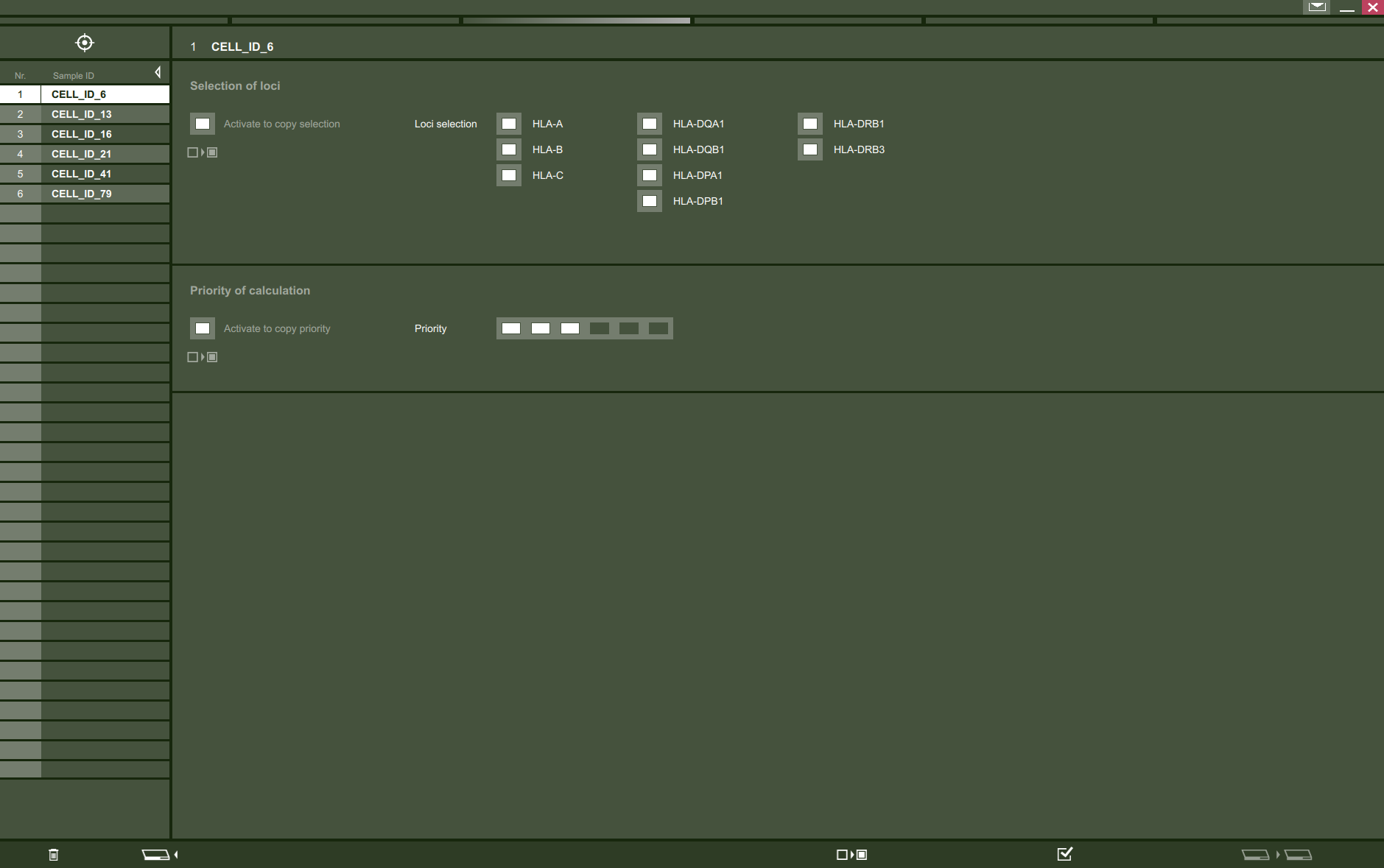

Figure 6. **Workflow Step 3**, locus selection and priorities. In this step, the user can select the HLA loci that will be analyzed and assign sample priorities. When moving to the next step, a higher prioritized sample is queued before samples with lower priority, when pushed to the calculation step.

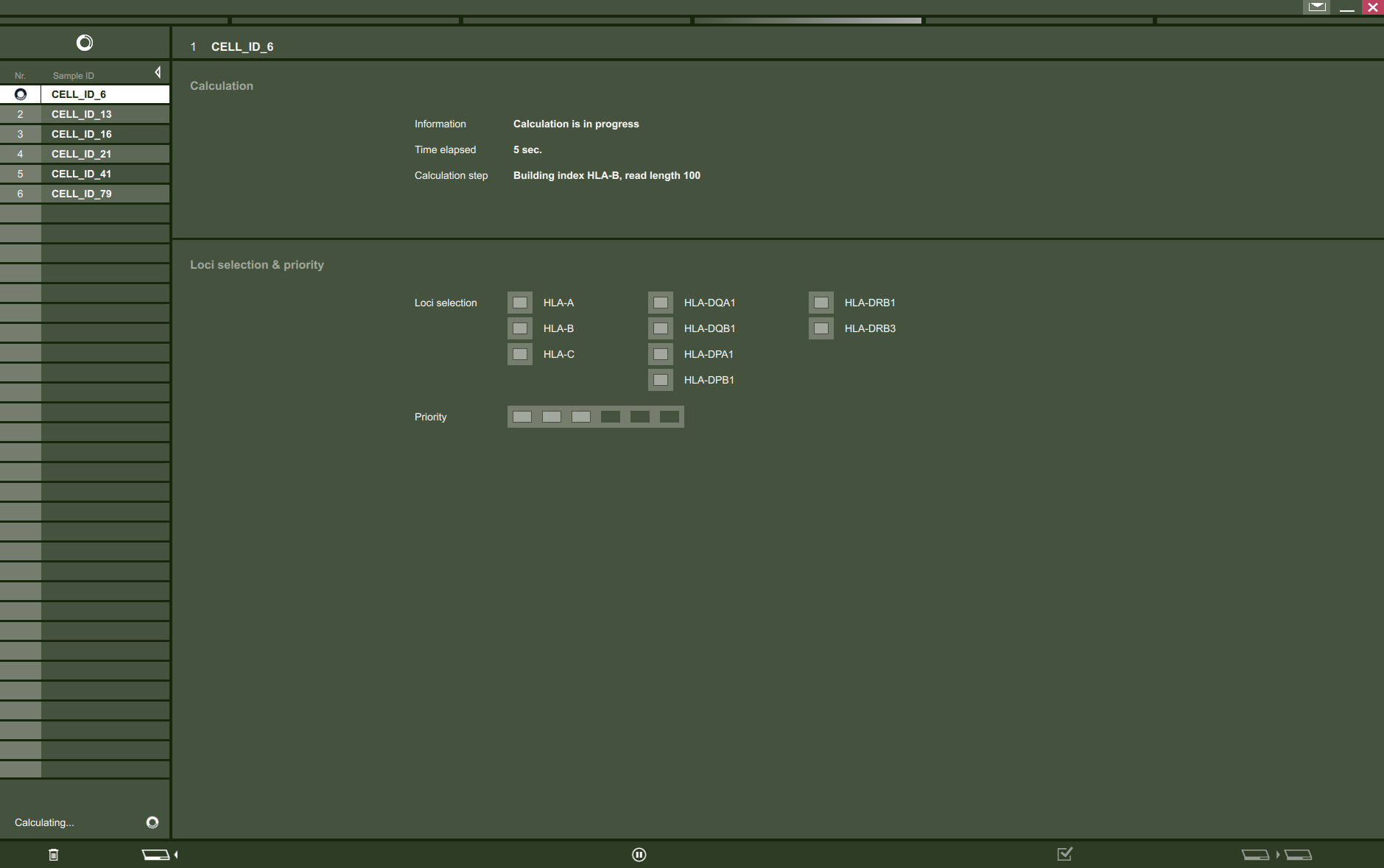

Figure 7. **Workflow Step 4**, calculation. Here is the queue of the samples that have to be analyzed. The current calculation step is shown at the top and the samples priority is shown below. The user can pause and continue the calculation.

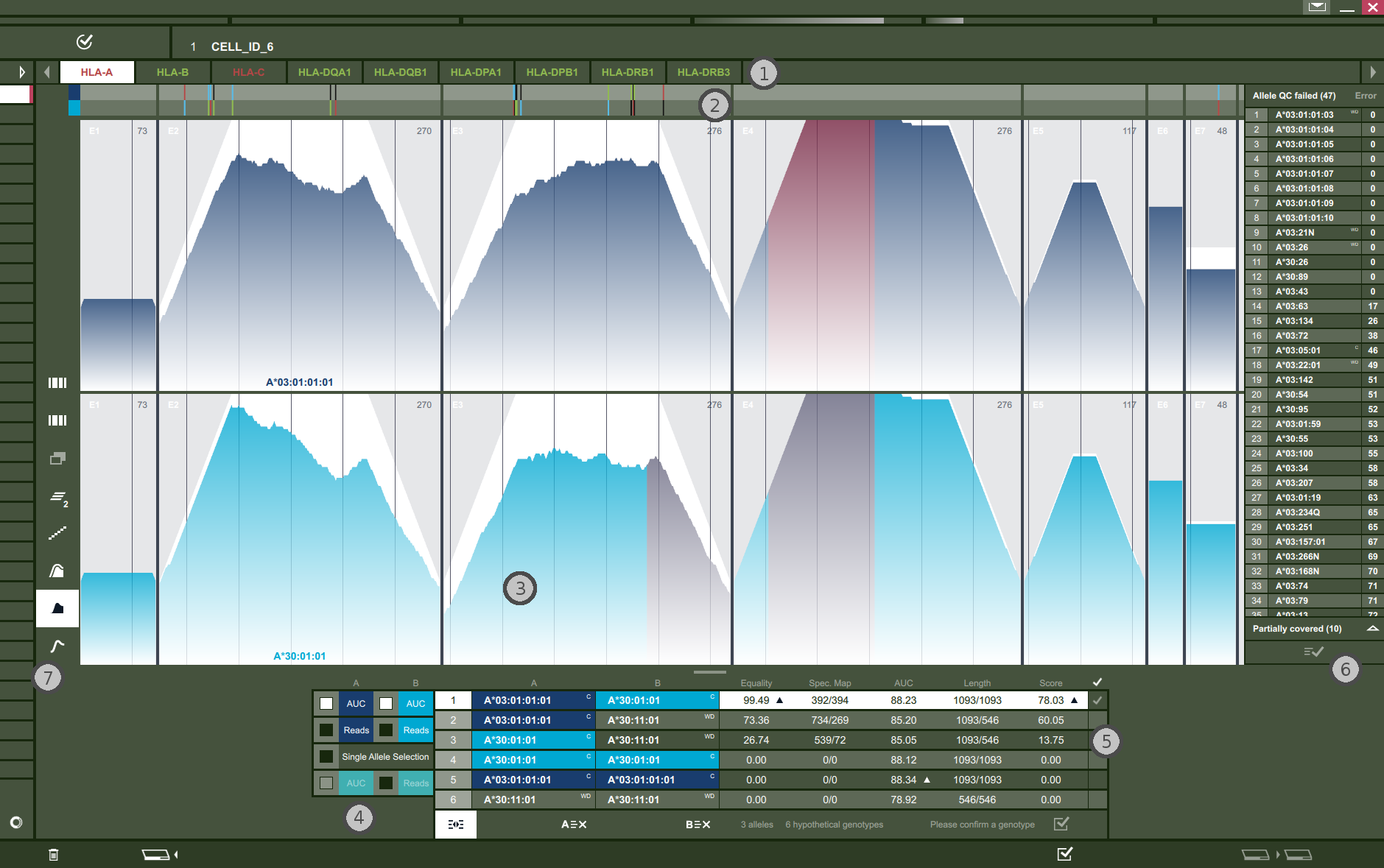

Figure 8. **Workflow Step 5**, validation. In this step, the user can evaluate the automated calling results. To perform an accurate validation it is important to understand the characteristics of the underlying data. The most important fact is the random fragmentation, that is normally performed for all samples prior to sequencing. Exploiting this knowledge, we expect an even distribution of the mapped reads across the entire locus and both alleles of a given genotype. The more balanced these distributions are, the more confident the calling of the genotype will be. The different sections are tagged by numbers. The different sections of the shown window are described in the following: **(1)** Contains the tabs for the different loci. If a locus is selected, the tab is highlighted (HLA-A) and the corresponding raw data and results are shown below. **(2)** Shows the consensus bar and all variations of the currently selected genotype (highlighted row in the genotype table (5)) as white bars. If clicked, which is the case in this figure, the consensus bar extends its height and separate color-coded bars show the different SNPs of the underlying alleles (A: green, C: blue, G: black, T: red, InDel: white). Here we show the extended version of the consensus bar. The raw data for each allele of a genotype is shown in section **(3)**. The default view is the exon-wise unique start point coverage. In an optimal alignment the coverage looks very similar to the white background that you can find at every exon (for HLA-A exons 1,4, 5 and 6). The here shown coverages at exon 2 and 3 are representing a typical coverage of good quality data and reflects what is normally observed. A red overlay signals sequence parts of the alleles that are identical to sequence parts of detected alleles in other loci (exon 4 and 5 and the 5’ part of exon 3 in allele A*30:01:01). This information may help to decide about false positives in some cases. The check boxes in section **(4)** are for toggling different raw data views. “AUC” shows the coverage while “Reads” show the underlying read mapping. If “Single allele Selection” is checked, the user can select a single allele and its raw data is shown in addition to the already selected alleles. The table at section **(5)** shows the ordered genotypes and it is sorted by the penalty (formerly score) coming from the automated calling. All possible genotypes of the alleles that passed the first filtering step are shown and ranked/scored here. The most important metrics are also shown. We refer to the **Supplementary Material** of the original publication^6^ for a detailed description of the different metrics. The buttons “A≡x” and “B≡x” are for deleting allele of column A or B of the selected genotype from the genotype table. The deleted allele moves to one of the tables of section 6. At section **(6)** the user can switch between two tables. The first table contains all alleles that are completely covered by reads but filtered out by the first filtering step (unbalanced read distribution). The second table lists some of the alleles that are not completely covered by mapped reads and is sorted by uncovered bases in ascending order. The number of entries can be determined in one of the initial dialogs on application start. Usually these alleles are only required for manual calling of very bad quality data and should only be performed by very experienced HLAssign users. Section **(7)** contains buttons to change the visualization style.

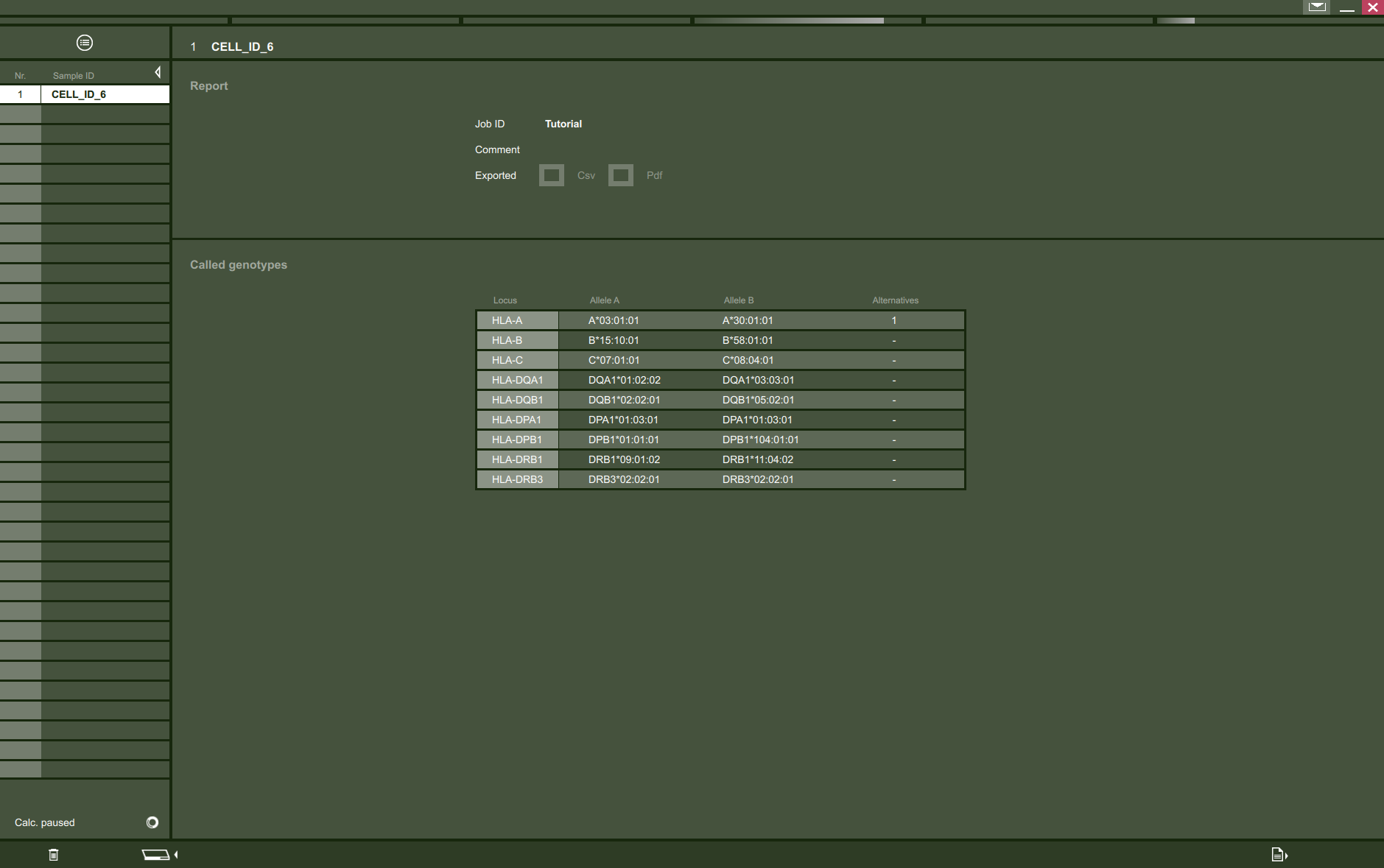

Figure 9. **Workflow Step 6**, report. The last step is the report and the user has a summarized view for each samples genotype calls. The results can be exported as a comma separated list for 1-n samples or as a PDF report for each sample separate.

**Validation**

The validation view (Figure 8) allows for validating the automated calling, removing alternatives and assigning the correct HLA calls. To perform a good validation it is important to understand the characteristics of the underlying data. The most important fact is the random fragmentation, that is performed for all samples prior to sequencing. Based on that knowledge we expect an even distribution of the mapped reads across the entire locus and both alleles of a given genotype. The more balanced these distributions are the more confident the called genotype is. Figure 10 shows an validation view screenshot, where the different sections are tagged by yellow numbers. The functions of the different sections are as follows:

Section **1)** contains the tabs for the different loci. If a locus is selected the tab is highlighted (HLA-A) and the corresponding raw data and results are shown below. Section **2)** is the consensus bar. It shows all variations of a given genotype as white bars. If clicked, the consensus bar extends its height and separate color coded bars show the different SNPs of the underlying alleles (A: green, C: blue, G: black, T: red, InDel: white). The raw data for each allele of a genotype is shown in section **3)**. The default view is the exone wise unique start point coverage. In an optimal data set the coverage looks very similar to the white background that you can find at every exone (e.g. exone 1,4, 5 and 6). The coverage of exone 2 and 3 is also a typical coverage of good quality data and reflects what you usually find. A red overlay signals sequence parts of the alleles that are similar to sequence parts of detected alleles in other loci. This information may help to decide about false positives in some cases. The check boxes in section **4)** are to toggle different raw data views. AUC shows the coverage while Reads show the underlying read mapping. If “Single allele Selection” is checked, the user can select a single allele and its raw data is shown in addition to the already selected alleles. The table at section **5)** shows the ordered genotypes and it is sorted by the score coming from the automated calling. All possible genotypes of the alleles that passed the first filter step are shown and ranked/scored here. The most important metrics are also shown. Please check the supplementary material of the original publication (doi: [10.1093/nar/gkv184](http://nar.oxfordjournals.org/content/early/2015/03/09/nar.gkv184.long) AddSuppFiles3) to understand their meaning. The first filtering, mentioned above, evaluates the read mappings. To pass the filter, all bases of an allele should be covered by mapped reads and all reads should map equally across all the exons. If unexpected gaps are detected, where no read mappings start, the sample is discarded (examples at Figure 11). This filtering step usually performs well if your samples have an AUC value > 85.0. Lower values introduce more uncertainty resulting in false positive HLA calls. Values lower than 50.0 require almost always manual validation of an experienced HLAssign user. The buttons “A≡x” and “B≡x” are for deleting allele of column A or B of the selected genotype from the genotype table. The deleted allele moves to one of the tables of section 6. At section **6)** the user can switch between two tables. The first table contains all alleles that are completely covered by reads but filtered out by the first filtering step. The second table lists some of the alleles that are not completely covered by mapped reads and is sorted by uncovered bases in ascending order. The number of entries can be determined in one of the initial dialogs on application start. Usually these alleles are only required for manual calling of very bad quality data and should only be performed by very experienced HLAssign users. As the size of this list influences memory usage and run time, keep the number low. Section **7)** contains buttons to change the visualization style.

Sometimes it is necessary to validate phasing information to get a high confident HLA call. Please read the next section “Phasing” to learn how to perform that with HLAssign.

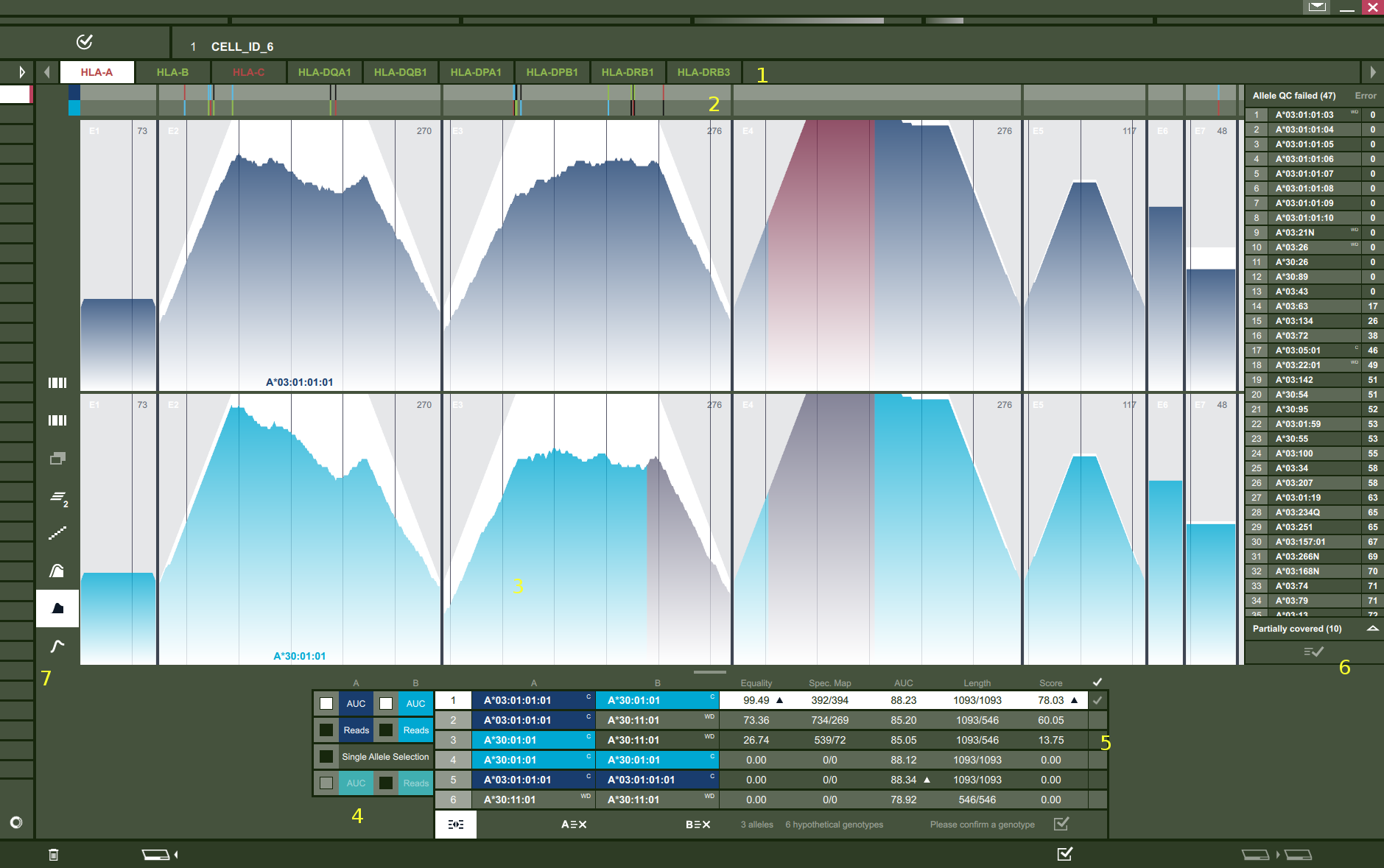

Figure 10. The validation view. Please read the section Validation for further details.

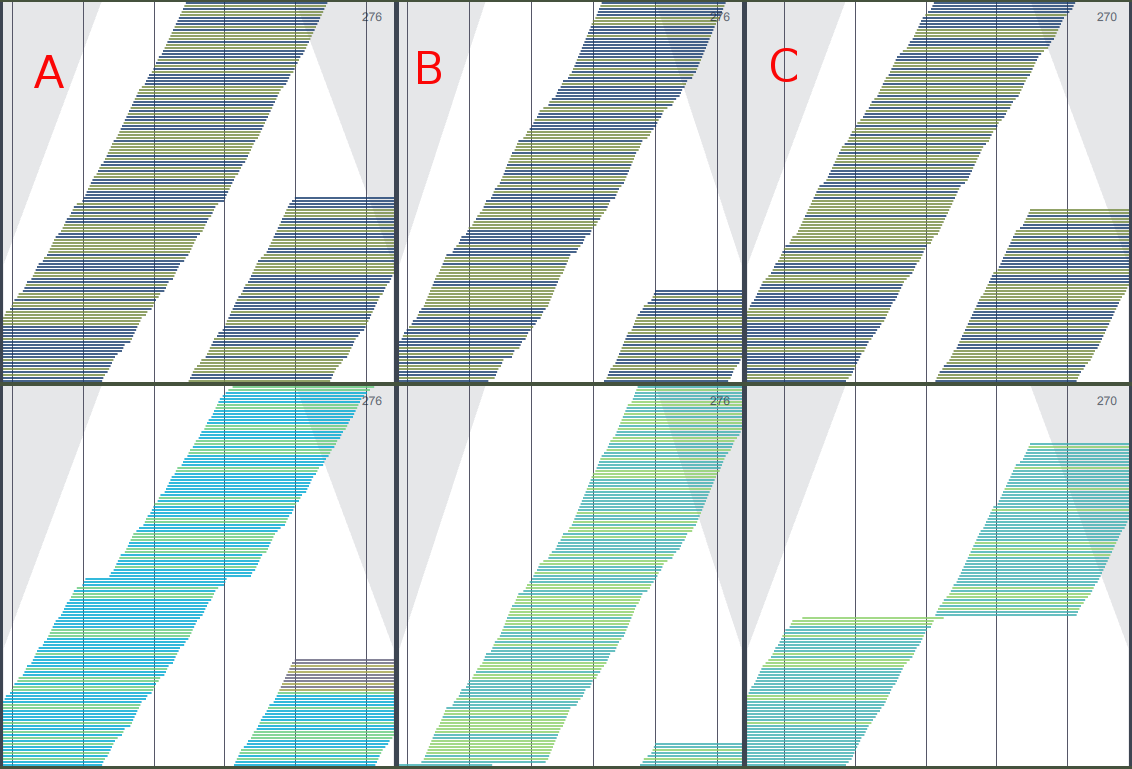

Figure 11. Examples of uneven read coverage. The lower panel of all three sections (A-C) shows examples of uneven read coverage across the exon. The example in section A shows a small gap where no read mapping starts (after the second light-grey vertical line). In section B the gap is at the 5’ part of the exone and a single read hides that phenomenon somehow. Although this is may be hard to detect it is obvious that the read mapping at the beginning of the exone is totally contrary to the rest of the exon. In section C is a huge gap in the middle of the exon.

**Phasing**

Phasing can be performed in the validation view by activating the Read check box for visualizing the read mappings. In that view all reads which have a second mate pair mapping have a yellow overlay (can be turned on/off by the uppermost button in section 7). If the user moves the mouse cursor over one of those reads, the read and its mate(s) are highlighted in yellow. The region covered by those reads is also highlighted at the bottom of the consensus bar. That makes it easier to identify all SNPs covered by that linked/phased reads. While pushing the left mouse button the before described feature is available for multiple read selection. Figure 12 and 13 illustrate how to identify false positives with the phasing functionality and how to assign the right HLA call HLA-B*15:01:01/18:02 in this example.

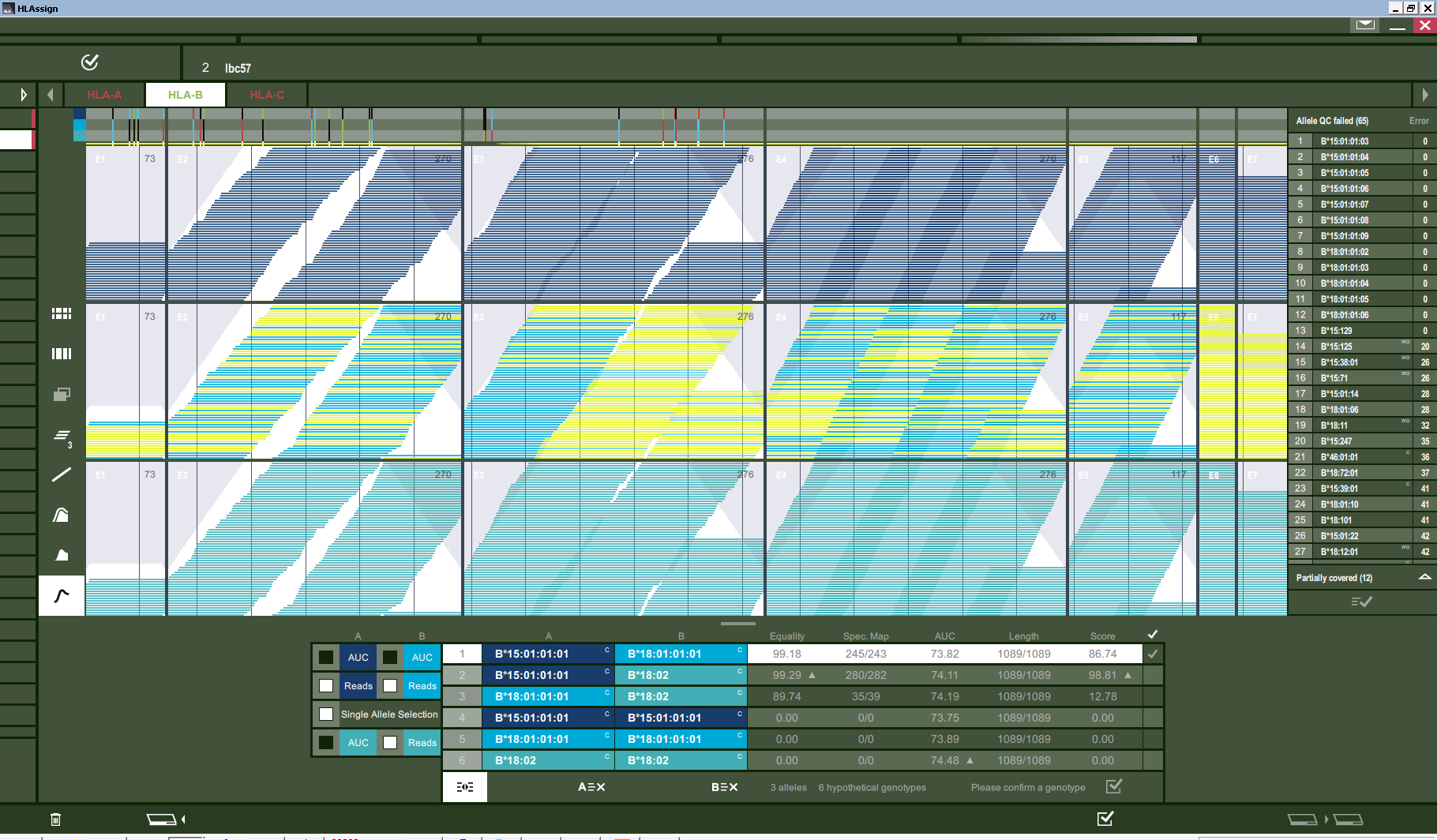

Figure 12. False positive allele in the genotype table. In this data view all paired end mappings of the allele in the middle (B*18:01:01:01) are highlighted. As a result all regions covered by those linked mappings are highlighted at the bottom of the consensus bar. It can be seen that there are no linked reads that allow phasing between the SNPs at the 5’ part of exone 3 and the rest of the exone 3 SNPs. Please take a look at Figure 13 to see how that phasing would look like for an true positive call.

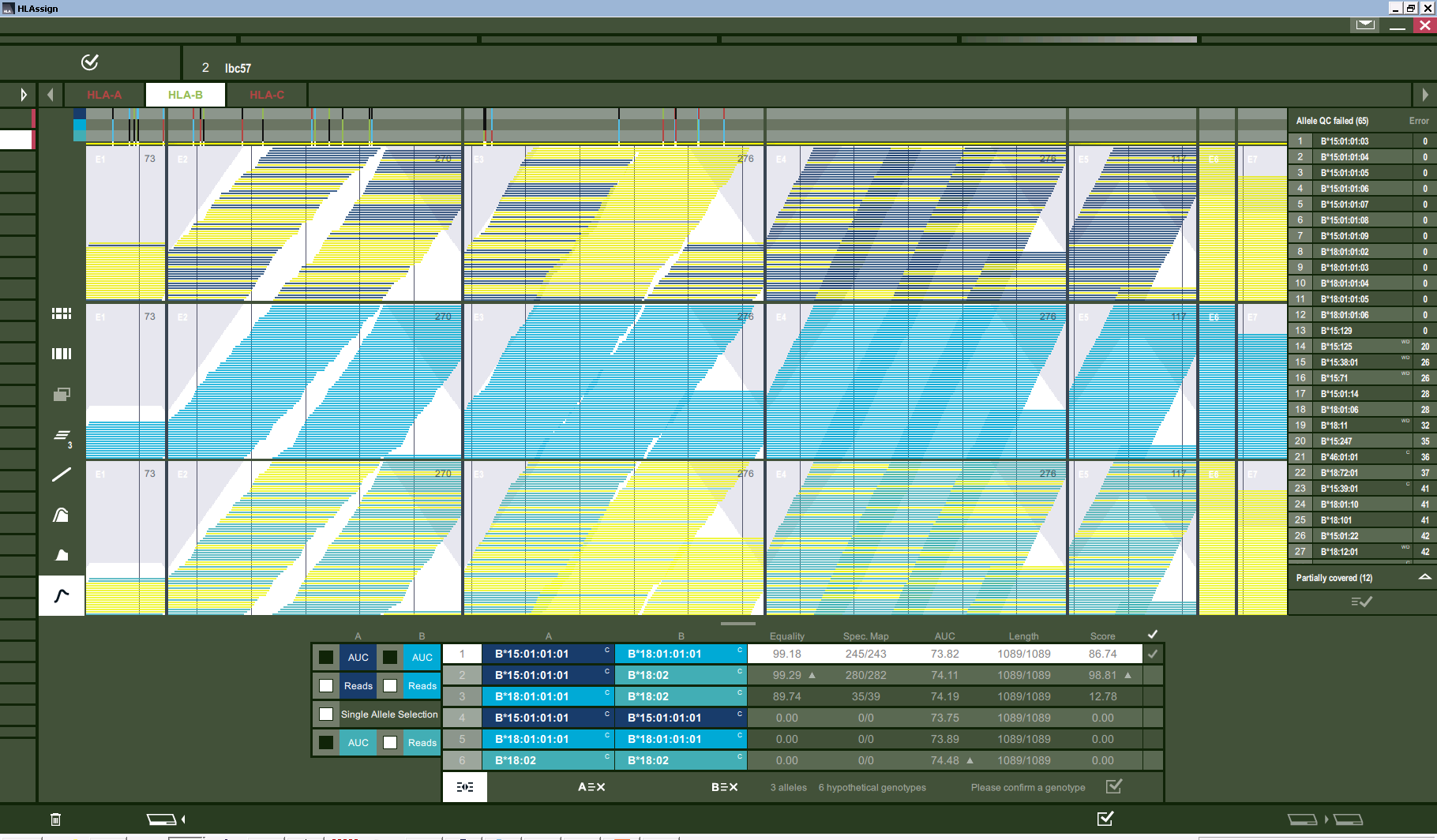

Figure 13. True positive allele in the genotype table. In this data view all paired end mappings of the alleles at the top and bottom (B*15:01:01:01 and 18:02) are highlighted. As a result all regions covered by those linked mappings are highlighted at the bottom of the consensus bar. It can be seen that all SNPs in every exone are somehow linked with each other. Sometimes, if there is a short intron, it is also possible to phase between neighbouring exones. Please take a look at Figure 12 to see how that phasing would look like for a false positive call.
