## Supplement S2, Examples data evaluation for "HLAssign 2.0: An advanced Graphical User Interface for the analysis of short and long read Human Leukocyte Antigen-typing data"

Typical phenomena that cause false positive signals can be evaluated visually and the functionality supports the user making the correct decisions in these cases. See Figure 1 and 2:

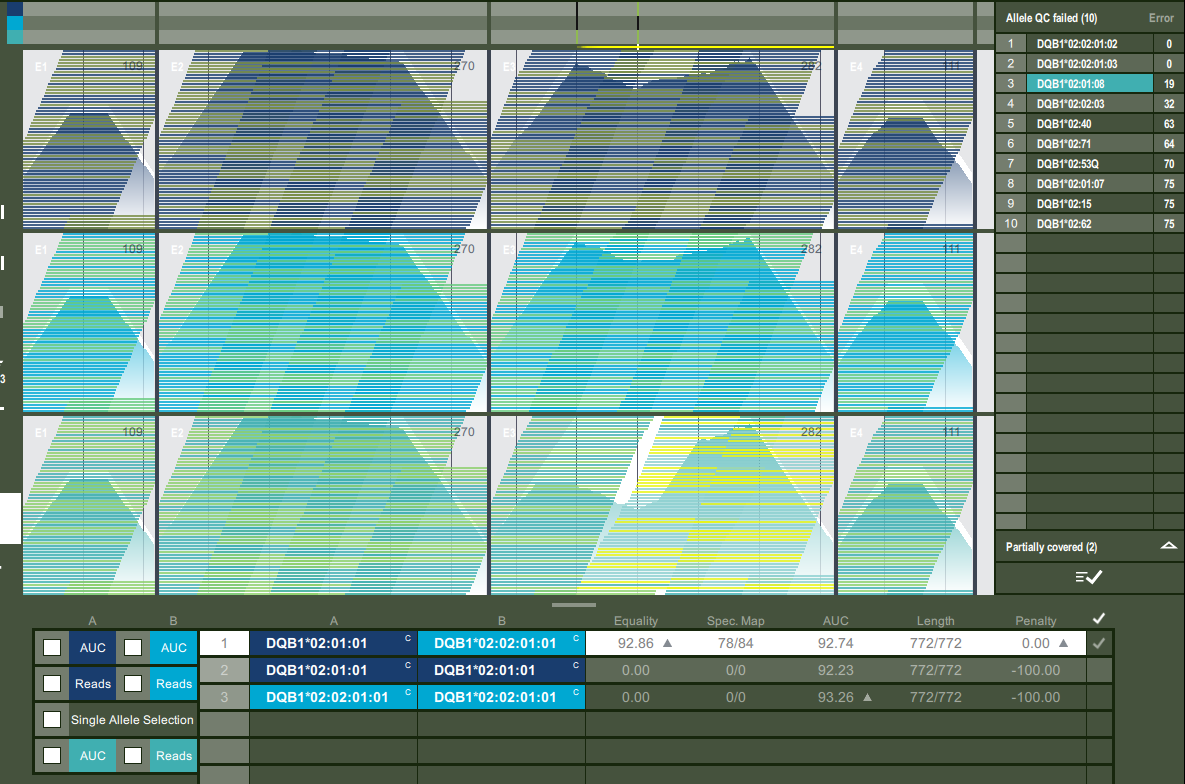

**Figure 1**: The DQB1 genotype is reported as 02:01:08 by STC-seq (lowest graph), but we can visualize that no read that maps downstream of cDNA position 354 has a linked paired read that maps to any position before. Or in other words, no haplotype with A-A in exon 3 can be found, but the two Haplotypes G-A (02:01:01) and G-G (02:02:01) can be found, so 02:01:01/02:02:01 is right here.

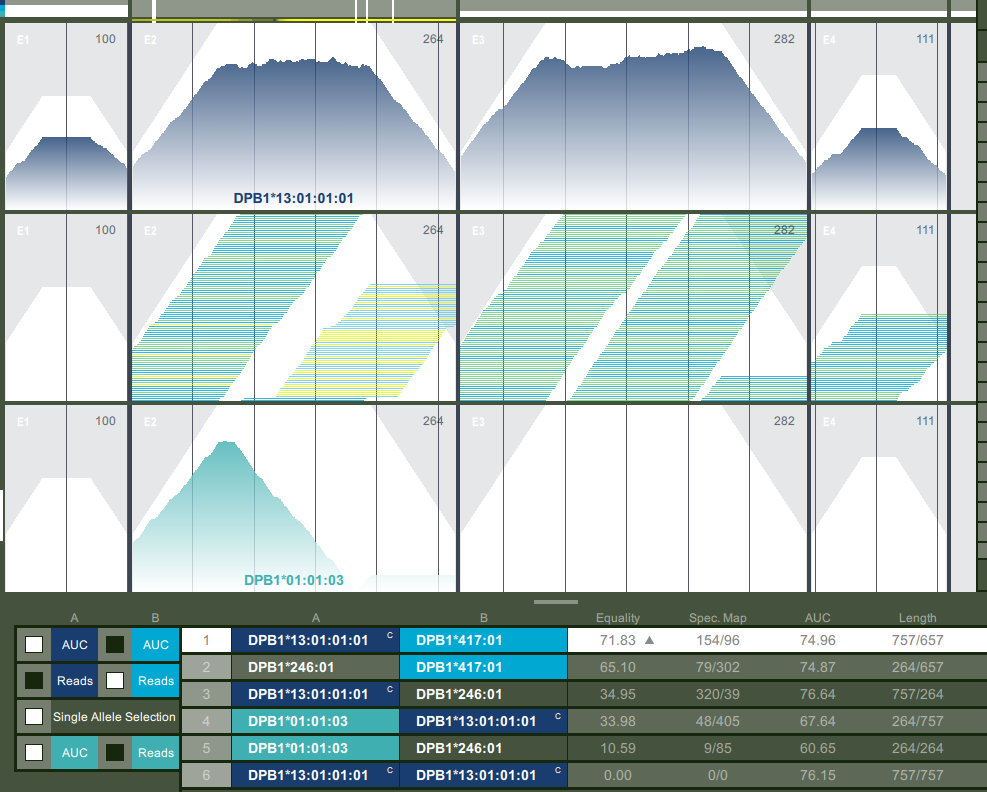

**Figure 2**: Another example for DPB1 where HLAssign reports 13:01:01/13:01:01, xHLA reports 01:01/13:01 and STC-seq reports 13:01:01/417:01. The figure shows that DPB1*01:01 is clearly negative and not completely covered by reads (a single nucleotide gap at exon 2). The allele DPB1*417:01 is harder to detect but visualization reveals that the reads overlapping the 3’ part of the exon 2 have a lot of paired ends mapping to all other regions of exon 2 except a single nucleotide close to the middle of exon 2.
